# Aβ Aggregates Bind the U1 Spliceosomal Ribonucleoprotein in Alzheimer Disease Brain

**DOI:** 10.64898/2026.06.02.729610

**Authors:** Youqi Tao, Anna E. Francis, Wen Liu, Christopher Watkins, David A. Bennett, Dennis J. Selkoe, Ryan A. Flynn, Andrew M. Stern

## Abstract

Missplicing due to U1 small nuclear ribonucleoprotein (U1 snRNP) insolubilization and dysfunction has been identified in Alzheimer disease (AD) brain. Cytoplasmic aggregation and mislocalization of the U1 snRNP partly co-localizes with tau neurofibrillary tangles, and some evidence for tau-U1 binding has been identified. However, tau-U1 co-localization by immunohistochemistry is only partial, and insoluble U1 small nuclear RNA (snRNA) binding proteins correlate better with Aβ than with tau in unbiased proteomics. While investigating the sedimentation characteristics of Aβ aggregates capable of diffusing out of AD brain tissue, we unexpectedly found that some were bound to small RNA. Aβ immunoprecipitation and deep sequencing revealed the U1 snRNA, with specific binding confirmed through reverse immunoprecipitation and oligonucleotide hybridization. Double immunoelectron microscopy revealed decoration of Aβ fibrils, more than tau fibrils, with the U1-70k protein, a component of the U1 snRNP. Immunofluorescence of unfixed cryostat sections but not formalin fixed paraffin embedded sections of AD brain revealed labeling of a subset of amyloid plaques with anti-U1-70k antibodies, confirmed by RNA *in situ* hybridization of the U1 snRNA. We conclude that a subset of AD brain Aβ aggregates are bound to the U1 snRNP at the edges of some amyloid plaques, explaining prior proteomics findings. These data provide a potential link between Aβ aggregation and spliceosome dysfunction and unite Aβ with other fibril-forming proteins across the neurodegenerative diseases whose aggregation is affected by RNA binding.

## Introduction

Alzheimer disease (AD) pathology is defined by insoluble, fibrillar deposits of extracellular amyloid-β (amyloid plaques) and intracellular tau (neurofibrillary tangles). The corresponding clinical syndrome involves multidomain cognitive impairment, most commonly with predominant memory loss. Changes to the expression of many genes and proteins differ between patients with Alzheimer disease and controls and may contribute to the phenotypic expression of plaques and tangles into cognitive impairment, ranging from inflammation, cellular energetics, RNA splicing, and others. Modern “omics” techniques have allowed the enumeration of these dysregulated pathways. Among them are proteins that are rendered insoluble in AD tissue, most famously Aβ and tau themselves. Components of the U1 spliceosomal small nuclear ribonucleoprotein (U1 snRNP), in particular the U1-70k protein, are also among the most insolubilized in AD brain compared to healthy control^1^. Loss of function of U1 snRNP due to insolubilization may lead to messenger RNA missplicing and cryptic polyadenylation^2^. The U1-70k protein possesses an intrinsically disordered region with alternating basic and acidic residues; this domain can cause aggregation, but amyloid fibrils of the U1-70k protein have not been observed^3,4^. Immunohistochemistry for U1 snRNP components in formalin-fixed tissue has revealed tangle-like cytoplasmic inclusions which sometimes overlap with neurofibrillary tangles, but sometimes do not, with some immunogold electron microscopy demonstrating evidence of anti-U1-70k co-localizing with paired helical filament tau, the components of neurofibrillary tangles^2,5,6^. Amyloid plaques have not been found to stain with anti-U1 antibodies in formalin-fixed sections. However, some inconsistencies in these data have emerged: the abundance of U1-70k in insoluble AD extracts correlates exquisitely with Aβ but only modestly with tau, and U1-70k aggregation appears in asymptomatic AD cases without significant tau aggregation^1^. This makes U1-70k aggregation a possible intermediary between Aβ aggregation and tau aggregation, and may also imply Aβ-associated, rather than tau-associated, U1-70k aggregation.

Inspired by literature demonstrating that aqueously extractable Aβ from human brain correlates with cognitive symptoms^7–9^, including observable *in vitro* neurotoxicity, we previously analyzed diffusible Aβ aggregates from human brain derived by soaking postmortem grey matter in aqueous buffer, followed by centrifugation^10^. In these “soaking extracts,”^11^ in which small and diffusible particles are extracted from the brain with as little chemical or mechanical disruption as possible, all detectable Aβ aggregates could be pelleted by ultracentrifugation^10^. Present among these diffusible, but still insoluble (particulate) Aβ aggregates, were amyloid fibrils with the same atomic structures as those present in the traditional sarkosyl-insoluble fraction^12^. This led us to the conclusion that toxic, aqueously extractable Aβ from human AD brain is diffusible but particulate and may include classically structured but short amyloid fibrils. We thus became interested in identifying binding partners of aqueously diffusible but particulate Aβ aggregates, including short amyloid fibrils, in human AD brain. Here, we report an unexpected direct interaction between Aβ fibrils and the U1 snRNP. We explain the lack of previous observation of this complex by the reliance on FFPE tissue sections. Our results are consistent with previous proteomic observations and suggest that an Aβ-U1 interaction may be an early event in AD pathogenesis.

## Results

### Diffusible Aβ aggregates in human brain are bound to RNA

Initially hypothesizing that some naturally occurring, diffusible Aβ aggregates would be bound to non-proteinaceous particles in the brain, especially lipoproteins, we subjected AD brain soaking extracts to isopycnic cesium chloride centrifugation, which uses overnight high-speed centrifugation in high-density CsCl solution to equilibrate particles on the basis of density rather than mass. Aβ in soaking extracts in CsCl gradients formed a broader and, to our surprise, denser peak than synthetic Aβ fibrils, as opposed to a less dense fraction than synthetic Aβ fibrils as would be hypothesized if lipids were bound (Fig 1A). The increase in density suggested the presence of nucleic acids, which generally exhibit greater density than proteins. Treatment of the extracts with RNase A shifted the soaking extract Aβ peak to a less-dense fraction, consistent with the presence of RNA in the Aβ particles (Fig 1B). RNase digestion in cellular and tissue lysates has been shown to cause the aggregation of both canonical RNA binding proteins and non-canonical RNA binding proteins, including Aβ^13^. We examined for this effect in soaking extracts by the more commonly used rate-zonal sucrose gradient centrifugation (sometimes confusingly called “density gradient centrifugation”), which uses a short centrifugation time on a less-dense gradient to separate particles based on their mass, but not density. RNase A digestion led to a shift in Aβ mass toward heavier fractions (Fig 1C), replicating prior findings and suggesting further that diffusible Aβ may be present in particles containing RNA.

**Figure 1.**
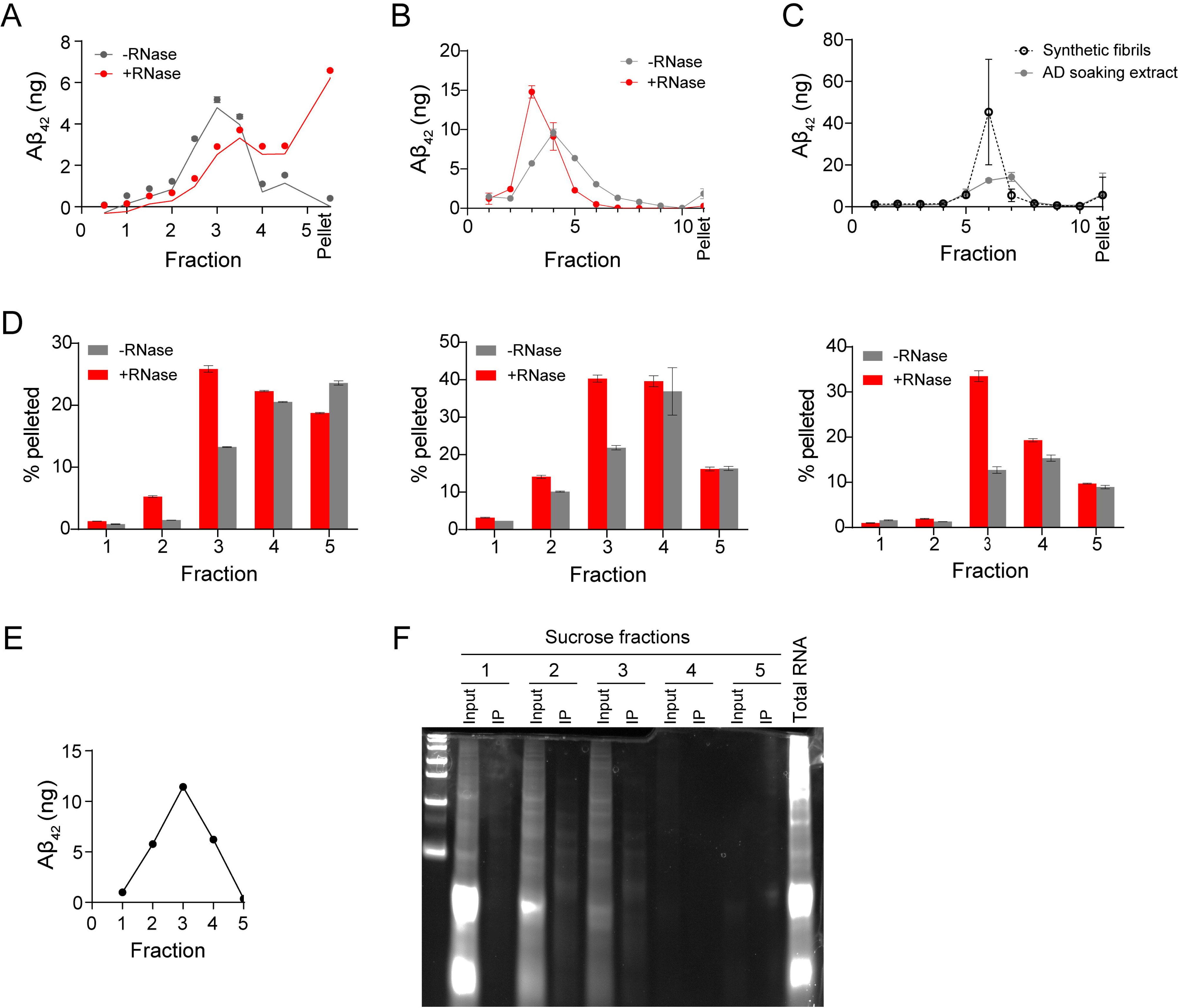
Some. **A**β **aggregates in AD soaking extracts are bound to RNA. (A)** Isopycnic CsCl gradients separate particles by density. Aβ_42_ in a soaking extract of AD2 equilibrated to a denser peak than fibrils produced from recombinant Aβ_42_, suggesting nucleic acids may be present. **(B)** Digestion of AD8 soaking extract with RNAse A shifted the Aβ_42_ to a less-dense peak, implying RNA as a component of Aβ aggregates. **(C)** Rate-zonal sucrose gradients separate particles by mass. RNAse A digestion of AD3 soaking extract prior to sucrose gradient centrifugation shifted Aβ_42_ to heavier fractions, consistent with the behavior of many RNA-binding proteins. **(D)** Sucrose gradient fractionation followed by RNAse digestion and re-centrifugation revealed preferential RNAse sensitivity in the lighter Aβ_42_-containing fractions (fractions 2-3) across three AD soaking extracts. **(E)** Comparable to crude preparations, sucrose gradient centrifugation followed by immunoprecipitation with Aβ-aggregate preferring antibody h1C22 revealed a peak around fraction 3. **(F)** Urea-PAGE of AD extracts separated by sucrose gradient centrifugation followed by h1C22 immunoprecipitation revealed that sucrose gradient fractions possess small RNAs preferentially in lighter fractions (“input”), but h1C22 IP isolated a distinct banding pattern of RNAs, particularly in fractions 2 and 3.

### Relatively lower molecular weight Aβ aggregates are bound to small RNA

Having established a possible complex of Aβ and RNA, we further used rate-zonal sucrose gradient centrifugation to determine if RNA-associated Aβ particles were lighter or heavier than non-RNA-associated Aβ. AD brain soaking extracts were first centrifuged on sucrose gradients, followed by RNase digestion, the reverse order of Fig 1C, and then the digests were further pelleted in a benchtop centrifuge. RNase A digestion caused increased pelleting of relatively lighter fractions 2-3 rather than heavier fractions 4-5 (Fig 1D), suggesting that relatively lighter Aβ aggregates may be RNA-associated. To begin to identify the RNA species, we performed sucrose gradient fractionation followed by immunoprecipitation with aggregate-preferring anti-Aβ antibody h1C22. As seen without immunoprecipitation, the Aβ peaked in the middle of the gradient around fraction 3 (Fig 1E). We extracted RNA from the crude sucrose fractions and after h1C22 immunoprecipitation and found that most of the RNA observed was small (<1,000 nucleotides) (Fig 1F). Most small RNA present in the soaking extracts fractionated in the top (lightest) fraction, where no Aβ was present, but a distinct banding pattern was resolved using urea-polyacrylamide gel electrophoresis after Aβ immunoprecipitation, particularly for the Aβ-containing fractions 2 and 3. These results altogether suggested that relatively lighter Aβ aggregates (fractions 2-3) in human AD brain are associated with a distinct pool of small RNAs.

#### Diffusible Aβ is bound to the U1 spliceosomal RNP

To identify the Aβ-associated RNA, we prepared soaking extracts from the angular gyrus of ten brains (five pathologic AD, five control) from the Religious Orders Study and Rush Memory Aging Project (ROSMAP) cohorts (Table 1). We subjected the ten extracts to sucrose gradient centrifugation followed by h1C22 IP from fraction 3, which contained the putative Aβ-RNA complex, followed by RNA sequencing (Fig 2A). We also sequenced the total input RNA for comparison. We determined a fold enrichment of RNA of sucrose gradient fractionation/h1C22 IP compared to input RNA separately for both AD and control cases (Fig 2B). We observed 660 RNAs that were differentially enriched by sucrose gradient fractionation/h1C22 IP in AD but not control. To identify biologically meaningful hits, we selected RNAs with 1) high expression (BaseMean > 20), 2) statistical enrichment in AD, and 3) higher AD enrichment compared to controls (non-AD samples). By these criteria, we observed a total of 16 RNAs (Table S2), of which the U1 small nuclear RNA (snRNA) was by far the most abundant. We observed some enrichment of other spliceosomal RNAs by the combined sucrose gradient/immunoprecipitation method across the full cohort, but only U1 demonstrated a preference for enrichment in AD (Fig 2C).

**Figure 2.**
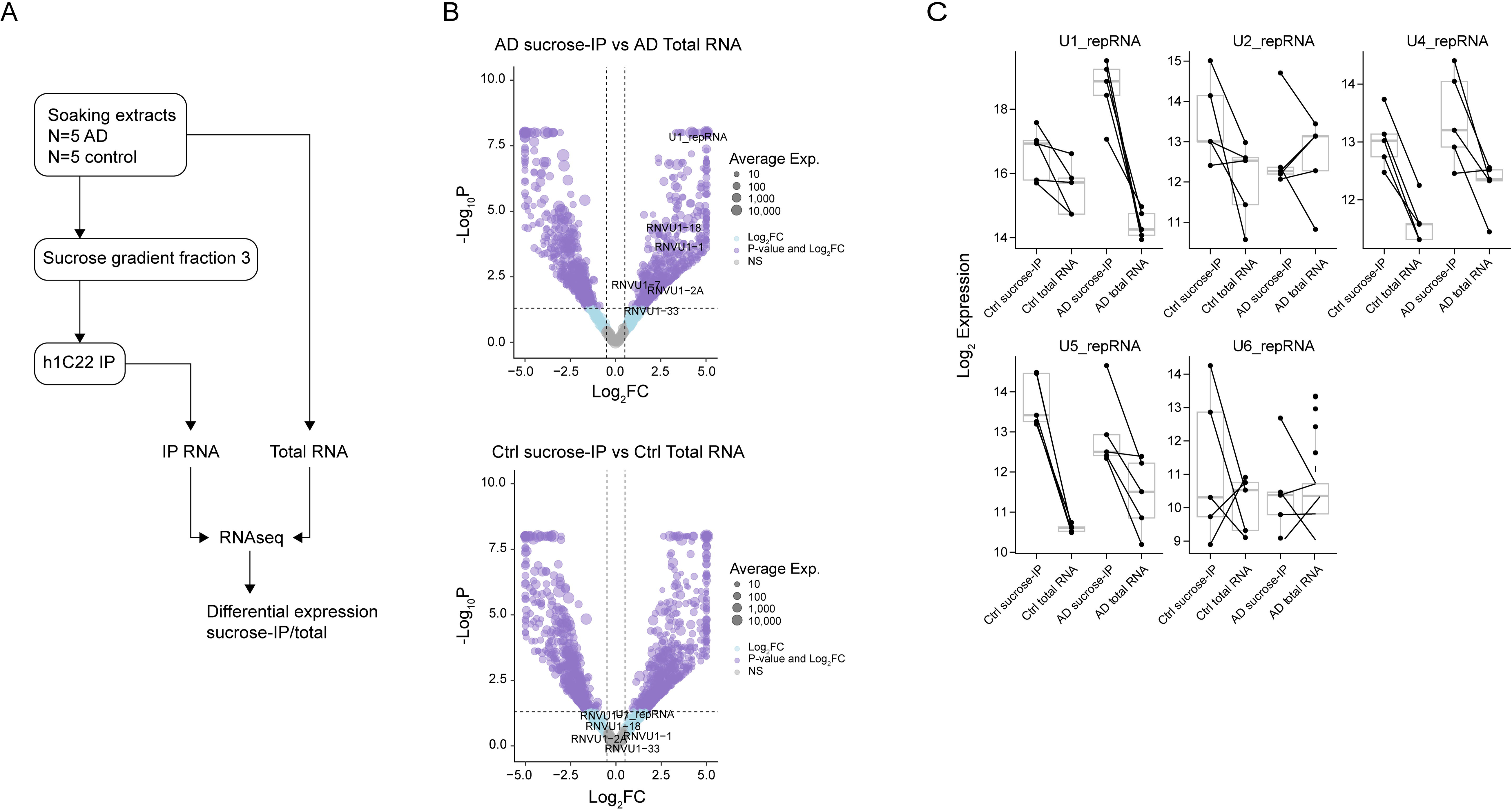
**A**β **binds the U1 snRNA. (A)** Schematic of enrichment for Aβ-associated RNA. Soaking extracts (N=10) were first separated by sucrose gradient fractionation, then fraction 3 was retained, buffer exchanged into TBS, and immunoprecipitated with h1C22 and extracted with Trizol. The RNA was sequenced and differential expression calculate compared to total input RNA separately for AD and control cases. **(B)** Volcano plots demonstrating specific enrichment of reads mapping to U1 snRNA and its variants in AD but not control tissue. Each dot is one gene and its size reflects the log_2_ of mean expression across the sample. **(C)** Box plots for each canonical spliceosomal snRNA. Only U1 shows specific enrichment by the sucrose-IP method in AD but not healthy control.

**Table 1.** ROSMAP cases used for sRNA sequencing. *By semiquantitative silver staining in angular gyrus.

| Case | Age | Sex | Braak | Aβ<sub>42</sub> (ng/g) in soaking extract | APOE genotype | Diffuse plaque score* | Neuritic plaque score* | Neurofibrillary tangle score* |
| --- | --- | --- | --- | --- | --- | --- | --- | --- |
| RAD1 | 84 | M | 5 | 153.2 | ε3/ε3 | 6 | 13 | 3 |
| RAD2 | 86 | F | 5 | 142.2 | ε3/ε4 | 12 | 5 | 15 |
| RAD3 | 88 | F | 5 | 152.5 | ε3/ε3 | 15 | 41 | 11 |
| RAD4 | 86 | M | 5 | 149.9 | ε3/ε3 | 10 | 30 | 6 |
| RAD5 | 89 + | F | 5 | 169.2 | ε3/ε3 | 20 | 14 | 5 |
| RCtrl1 | 83 | M | 2 | 0.4 | ε3/ε4 | 0 | 0 | 0 |
| RCtrl2 | 89 + | F | 1 | 0.5 | ε3/ε3 | 0 | 0 | 0 |
| RCtrl3 | 89 + | F | 3 | 27.9 | ε3/ε4 | 0 | 0 | 0 |
| RCtrl4 | 82 | M | 3 | 1.4 | ε3/ε3 | 0 | 0 | 0 |
| RCtrl5 | 89 + | F | 4 | 1.6 | ε3/ε4 | 0 | 0 | 0 |

We next sought to validate our evidence for a direct interaction between Aβ aggregates and the U1 snRNP. We first confirmed the RNA-seq by RT-qPCR, finding that h1C22 preferentially enriched U1 snRNA but not U2 from AD brain (Fig 3A). We also found that h1C22 co-immunoprecipitated the U1-binding protein U1-70k in AD brain but not control brain (Fig 3B). Immunoprecipitation with AT8, which binds phosphorylated tau, also immunoprecipitated U1-70k as has been reported^14^ but to a lesser degree than h1C22 (Fig 3C). In the reverse direction, UV crosslinking followed by hybridization to a biotinylated anti-U1 oligonucleotide and streptavidin pulldown could isolate Aβ in an RNase- and UV dose-dependent manner (Fig 3D). Immunoprecipitation of U1-70k also resulted in detection of Aβ, in particular Aβ_42_, which is more commonly found in AD brain aggregates, compared to Aβ_40_ (Fig 3E). Digestion with RNsse A did not disrupt the ability to IP Aβ with the anti-U1-70k antibody, suggesting that while the Aβ-U1 snRNP complex involves the U1 snRNA, the interaction does not strictly depend on the presence of the RNA.

**Figure 3.**
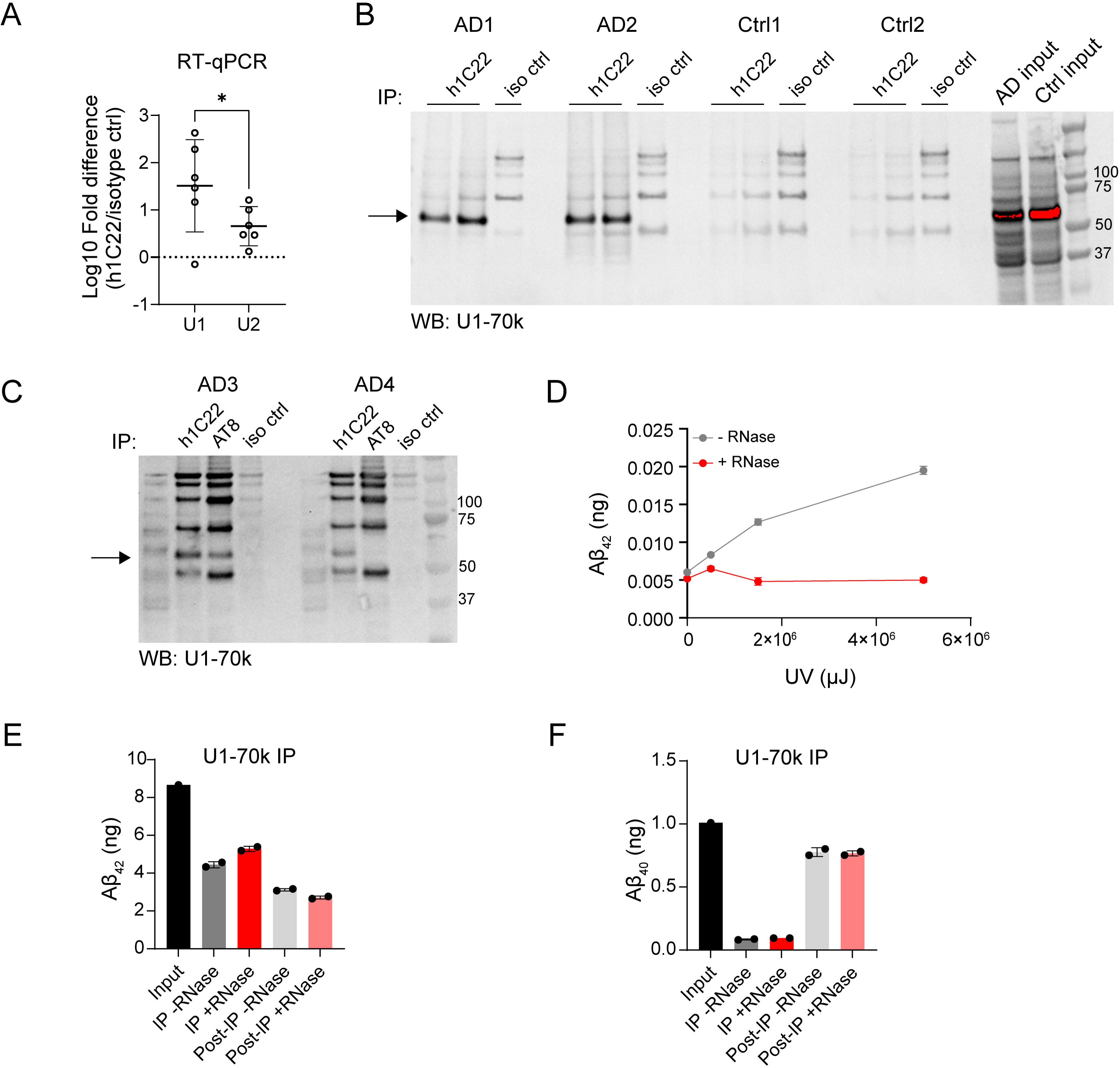
Validation of the. **A**β**-U1 snRNP complex. (A)** RT-qPCR of five AD soaking extracts (AD3, AD9, AD10, AD11, AD12) following IP with h1C22 or isotype control revealed enrichment of U1 more than U2 RNA. *P<0.05 by paired student’s t-test. **(B)** Co-IP with h1C22 followed by Western blot for the U1-70k protein revealed an interaction in AD brain but not control brain. **(C)** h1C22 could co-IP more U1-70k than anti-phosphotau antibody AT8. **(D)** UV crosslinking of AD4 soaking extract followed by hybridization to biotinylated anti-U1 oligonucleotide and pulldown with streptavidin revealed RNAse-dependent and UV-dependent isolation of Aβ_42_. **(E)** U1-70k IP from AD1 soaking extract revealed co-IP of ∼50% Aβ_42_ present but only ∼10% of Aβ_40_present; neither was affected by RNAse A digestion.

We next examined for U1-70k using immunogold labeling in soaking extracts (Fig 4). Using an anti-U1-70k antibody alone with protein A-nanogold, we found immunoreaction with thin fibrils that resembled Aβ. Adding an anti-Aβ antibody directly conjugated to larger gold particles, we observed frequent co-labeling of fibrils, suggesting a direct interaction between Aβ fibrils with U1-70k protein. Substituting a rabbit IgG control for the anti-Aβ antibody substantially reduced co-staining. Last, substituting an anti-phosphotau antibody immunolabeled tau fibrils, but we saw only rare co-localization with U1-70k, not appreciably above the background staining with the control rabbit IgG.

**Figure 4.**
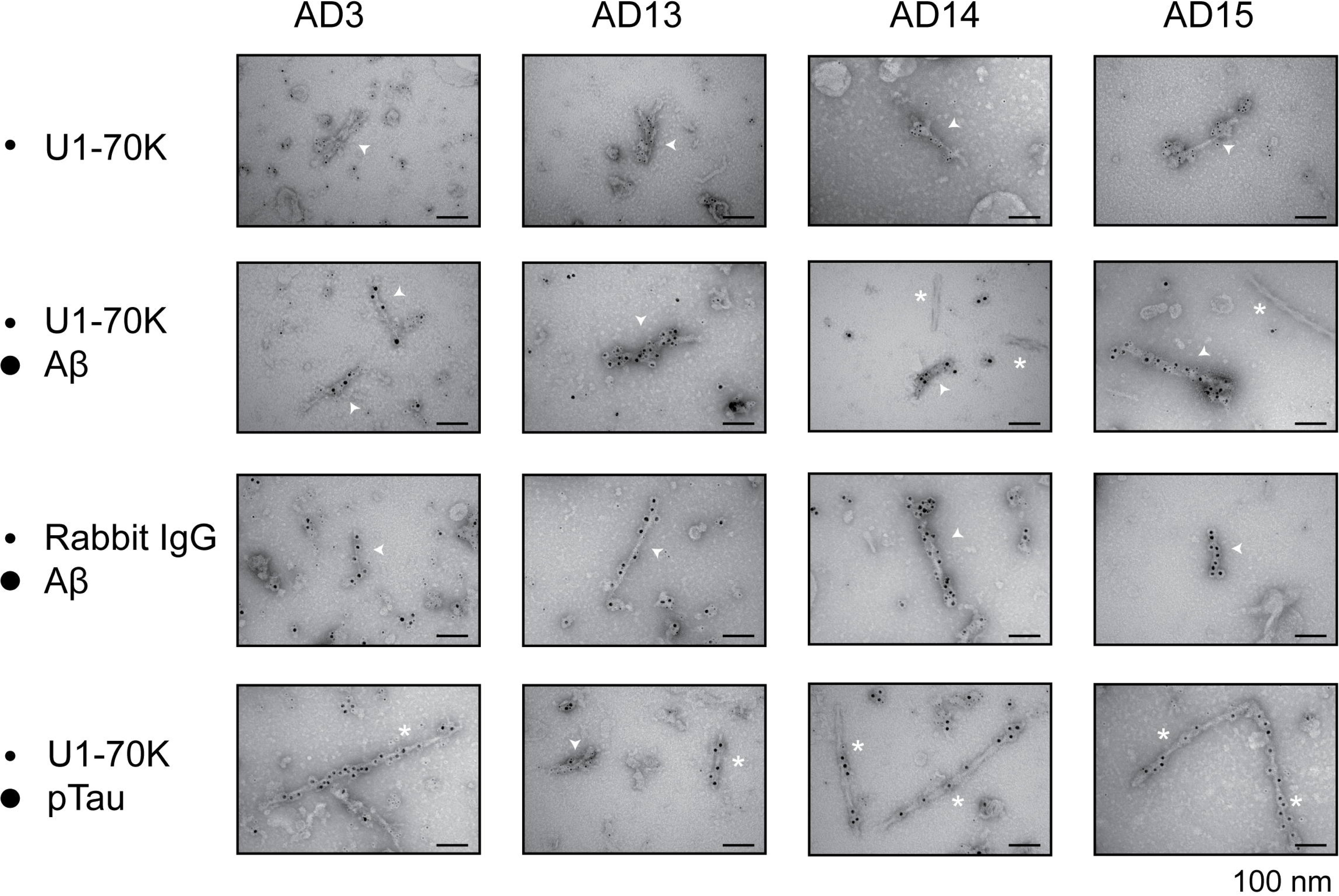
Double immunogold electron microscopy of the. **A**β**-U1 complex.** Figure 4**. Double immunogold electron microscopy of the A**β**-U1 complex.** Re-pelleted aqueous soaking extracts from four AD cases were sequentially labeled with anti-U1-70K (ab83306) followed by 5-nm protein A-gold, and either anti-Aβ (D54D2) or anti-phosphorylated Tau (AT8) directly conjugated to 10-nm gold particles. As a negative control for the U1-70k antibody, anti-rabbit IgG (DA1E) with 5-nm protein A-gold was used in combination with 10-nm gold-conjugated anti-Aβ (D54D2). Co-labelling revealed abundant Aβ fibrils associated with U1-70K (arrowheads). Tau fibrils were observed and only associated with infrequent U1-70k labeling (asterisks). Scale bar = 100 nm. Full-scale images are shown in Fig S4.

We conclude that individual Aβ fibrils bind directly to U1-70k, more than tau fibrils, consistent with our immunoprecipitation results and with prior proteomics literature^1^. We conclude that in the AD brain, the U1 snRNP forms a *bona fide* complex with Aβ aggregates.

#### The U1 snRNA interacts with Aβ at the amyloid plaque periphery

Previous immunohistochemistry and immunofluorescence studies have found that the U1-70k and other U1 binding proteins form intracellular tangle-like inclusions in neurons in AD brain^2,5,15^. The intracellular inclusions sometimes co-localized with tau but sometimes did not. Some plastic-embedded transmission immunogold electron microscopy revealed gold particles along putatively intracellular PHF tau^16^, the major constituent of tangles. These findings are consistent with our observation that AT8 immunoprecipitated U1-70k, albeit less than h1C22 immunoprecipitated, and our observation of occasional immunogold labeling of tau fibrils with anti-U1-70k antibodies. In previous studies, there was no staining of amyloid plaques by anti-U1-70k antibodies. We found the same: in formalin fixed paraffin embedded sections, we did not see labeling of amyloid plaques or their periphery by an anti-U1-70k antibody. However, we turned to unfixed cryostat sections recognizing that the fixation and antigen retrieval process can sometimes interfere with antibody binding. In unfixed parietal cortex, we found frequent co-staining of U1-70k within, and more often in the peripheral halo of, amyloid plaques (Fig 5A).

**Figure 5.**
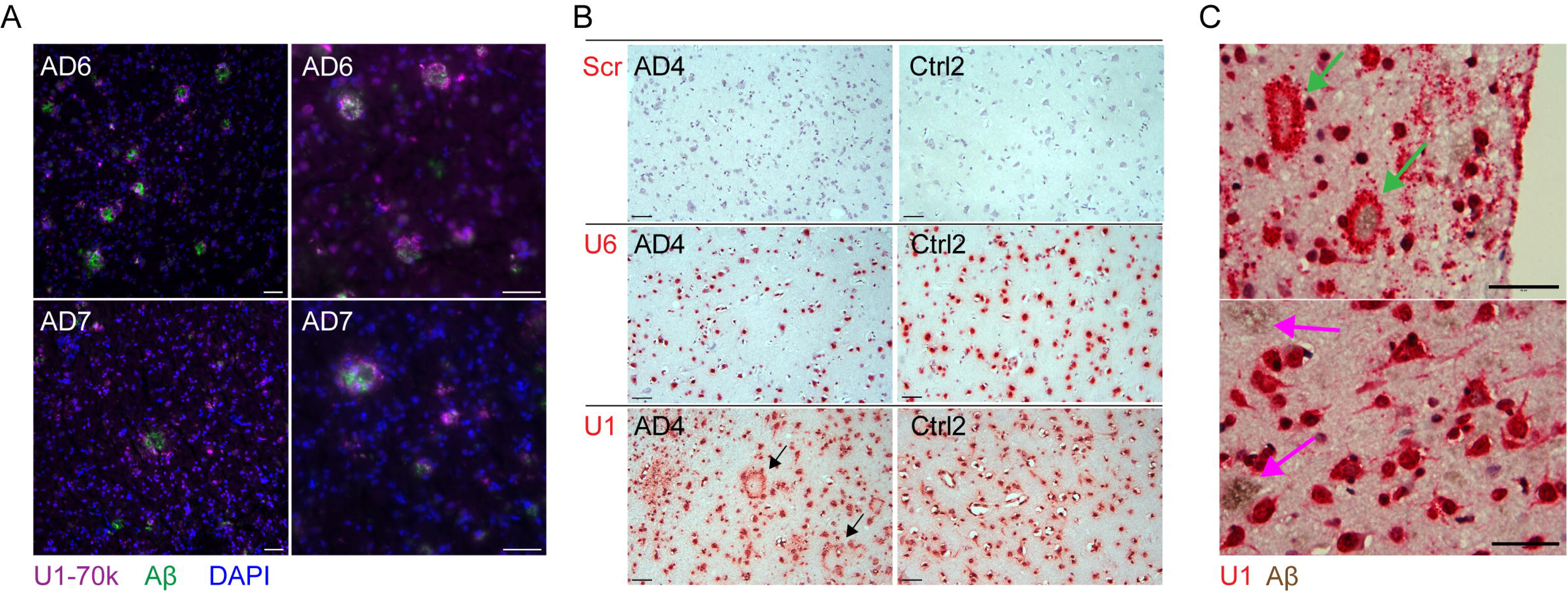
The U1 snRNA is present at some amyloid plaque edges. **(A)** Immunofluorescence amyloid plaques (6E10) and U1-70k in 50-µm cryostat sections from unfixed frozen AD brain revealed the presence of U1-70k around some plaques. **(B)** In FFPE tissue, monoplex RNA *in situ* hybridization to U1 snRNA revealed plaque-like structures (green arrows) in AD brain that were not present in control brain or when probing for the U6 snRNA or a scrambled probe. **(C)** Duplex RISH with Aβ immunohistochemistry (6E10) in brain AD4 revealed intense peri-plaque staining of U1 for some IHC-positive plaques (green arrows) but not for others (purple arrows). Scale bar = 50 µm.

Because available antibodies were unable to stain plaque-associated U1-70k in fixed tissue, we used RNA *in situ* hybridization (RISH). We found that probing for the U1 snRNA displayed intense plaque-sized rings in addition to the expected nuclear staining in AD (Fig 5B). The halos were not observed in control non-AD cases, nor were they observed with an anti-U6 snRNA probe; both of the latter only displayed expected nuclear staining. Co-staining with immunohistochemistry for Aβ simultaneously with RISH confirmed the presence of intense U1 snRNA co-localization at the periphery of plaques (Fig 5C). The intense U1 halo was present for only a subset of amyloid plaques, even within the same tissue section. We conclude that the U1 snRNP accumulates at the edges of a subset of amyloid plaques in Alzheimer disease brain.

## Discussion

This work has several limitations. First, it was performed entirely on postmortem human tissue. While this avoids error from model system deficiencies, our conclusions may be confounded by postmortem artifact – it is plausible that some Aβ-RNA associations may occur postmortem. The presence of U1 RNA and U1-70k at the edges of plaques in tissue sections, however, mitigates this risk.

Second, we have found substantial non-specific binding of anti-Aβ antibodies to material in brain extracts, including RNA and protein. This has been our experience for most aggregate-preferring Aβ antibodies tested at various stringencies. In one previous study, IP with the anti-Aβ antibody B24 depleted total protein from a brain extract by about 20%^17^. Others have found that lecanemab, an anti-Aβ antibody, binds to fibrinogen^18^. In this study, h1C22 could IP some RNA even in some control cases when no detectable Aβ aggregates were present. We suggest caution in interpreting studies in the literature positing the identification of Aβ binding proteins using IP without validation by antibody-independent methods. Because of this non-specific isolation of RNA even in control brain and low overall abundances, we were unable to identify, in this limited sample, small RNAs beyond the U1 snRNA that definitively Aβ-bound specifically in Alzheimer disease. In the future, we intend to use further non-antibody-based methods for Aβ enrichment plus include larger numbers of cases.

RNA has previously been described as a component of amyloid plaques^19,20^ but individual RNAs have not. In mouse models, nucleic acid-containing plaques were surrounded by reactive microglia expressing interferon signaling genes, whereas those lacking nucleic acids were not^21^. In this study, only a subset of amyloid plaques displayed U1 co-staining. With our limited sample size, it is not possible to know whether plaque morphology (*e.g.* diffuse, cored) or surrounding pathology relates to U1 co-staining, but this will be a goal of future studies. It is plausible that an Aβ-RNA complex is a particularly pro-inflammatory stimulus to microglia via toll-like receptors.

The mechanism of formation of the Aβ-U1 complex is mysterious and requires further study. Aβ, and especially fibrils thereof, is canonically extracellular, whereas the U1 snRNP is canonically intranuclear, where it performs pre-mRNA splicing. However, numerous small RNAs, including U1, have been reported in the extracellular space, often in fractions containing extracellular vesicles^22^. Glycosylated U1 snRNA and associated binding proteins including U1-70k have also been identified on the cell surface of cell lines^23^. Future studies will determine whether Aβ interacts with the U1 snRNP in the extracellular space, at the cell surface, or elsewhere, and what the effect of this interaction on brain cells is.

Despite key limitations, we feel convinced of a true complex between Aβ and the U1 snRNP. Far more work is required to understand in more detail how this complex forms, what other components of the complex are present, and what its functional or deleterious consequences might be in Alzheimer disease, if any.

## Methods and Materials

### Human brain tissue

Tissue for this study derived from one brain bank (BWH Neuropathology Tissue Repository and Clinical Data Bank) and one biorepository (Rush ADRC). For both sources, one hemisphere is fixed for histopathologic analyses, including those in this study, and one hemisphere is sliced fresh coronally and frozen in slices. Biochemical and histological analyses were performed on tissue from the BWH Neuropathology Tissue Repository and Clinical Data Bank from patients at Mass General Brigham Department of Neurology undergoing standard neuropathologic autopsy workup (Table S1). Soaking extract preparation (see below) proceeded using pooled grey matter from multiple regions of 1-2 coronal slices. *APOE* genotypes are presented when available. Tissue for small RNA sequencing was derived from frozen tissue from the angular gyrus (Brodmann areas 39/40) of cases from the Religious Orders Study and Rush Memory Aging Project (ROSMAP) cohorts of the Rush Alzheimer Disease Center (Table 1).

Both ROS and MAP enroll persons without dementia who agree to annual clinical evaluation and brain donation^24^. Both ROS and MAP studies were approved by an Institutional Review Board of Rush University Medical Center and all participants signed informed and repository consents and an Anatomic Gift Act. All cases undergo detailed histopathologic analyses including quantitative and semiquantitative scoring of amyloid plaques and neurofibrillary tangles.

### Human brain soaking extracts

Brain soaking extracts were performed as described previously^10,11^. Grey matter was dissected from fresh frozen cortex of individuals with AD and minced using a McIlwain tissue chopper (razor blade) set at 0.05 mm. The resultant brain bits were soaked for 30 min in a 1:5 ratio (w:v) of TBS extraction buffer (25 mM Tris supplemented with 150 mM NaCl, as well as protease inhibitors 5 μg/ml leupeptin, 5 μg/ml aprotinin, 2 μg/ml pepstatin, 120 μg/ml 4-benzenesulfonyl fluoride hydrochloride, and phosphatase inhibitor 5 mM NaF, pH 7.2) in 50-ml Eppendorf Protein LoBind tubes, and then spun at 2,000 *g* in a Fiberlite F14–14 × 50cy rotor in a Sorvall Lynx 6000 centrifuge for 10 min at 4°C (pelleting distance ∼4 cm depending on volume). TBS was chosen because its physiologic pH and salt concentration. We chose this over artificial CSF because the latter is not as well buffered. The top ∼90% of the supernatants were transferred to thin-wall polypropylene tubes (Beckman 331372) and spun using an SW41Ti rotor at 40,000 rpm for 110 minutes in an Optima L90K ultracentrifuge at 4°C (pelleting distance ∼9 cm). The top ∼90% supernatants were retained and frozen at −80°C in 1 ml aliquots.

### Aβ ELISA

ELISA quantitation of Aβ used in-house Meso Scale Discovery (MSD) platform assays described previously^10,17^. Briefly, samples containing GuHCl were diluted before ELISA such that the GuHCl concentration was <0.25 M, to avoid interference with the antibody reaction. All steps were at room temperature. Plates were coated with capture antibody in PBS overnight and blocked in 5% MSD Blocker A in TBS with 0.05% Tween-20 (TBS-T) for 1 h. Samples were then applied for 1.5 h after being diluted in 1% Blocker A in TBS-T. Plates were washed 3 times in TBS-T, then biotinylated detector antibody and MSD Strepavidin-Sulfotag (1:5,000) were applied for 1.5 h. After three more washes in TBS-T, the plates were detected with 2x MSD read buffer. The lower limit of quantification (LLoQ) was defined as the lowest standard with a luminescence value at least twice the blank average, and the lower limit of detection (LLoD) was defined as the lowest standard with values greater than the blank average plus twice the blank standard deviation.

### Isopycnic cesium chloride ultracentrifugation

Soaking extracts were thawed and optionally incubated with or without Monarch RNAse A (New England Biolabs) 20 µg/ml for 30 minutes at 37°C. Then, 0.1-0.25 ml extract was loaded onto a cesium chloride solution (1.4 g/cm^3^) in a thickwall polycarbonate tube (Beckman 343778) to a total volume of 1 ml. The tubes were centrifuged in a TLA120.2 rotor at 120,000 rpm overnight (∼19 hours) at 4°C to allow them to come to density equilibrium. Fractions (0.1 ml) were collected by pipetting from the top of the tube, then denaturing in 5 M GuHCl, including any pellet overnight at 4°C followed by ELISA.

### Rate-zonal sucrose gradient ultracentrifugation

Soaking extracts were thawed and optionally incubated with or without Monarch RNAse A (New England Biolabs) 20 µg/ml for 30 minutes at 37°C. Then, 0.95 ml extract was loaded on top of a sucrose gradient consisting of 1 ml 40% sucrose, 1 ml 25% sucrose, and 1.5 ml 5% sucrose in low-salt TBS (20 mM Tris-HCl, 150 mM NaCl, pH 7.4) in a thinwall polypropylene tube (Beckman 326819). The gradient was centrifuged at 55,000 rpm in an SW55Ti rotor for 30 minutes with maximum acceleration and deceleration at 4°C. Fractions were carefully pipetted off the top for downstream analysis. For ELISA, fractions were directly denatured using 5 M guanidine hydrochloride (GuHCl). For immunoprecipitation and/or small RNA sequencing, 0.35 ml soaking extract was diluted to 1 ml in low-salt TBS and then subjected to rate-zonal sucrose gradient ultracentrifugation as above. Fraction 3 was then buffer exchanged back into low-salt TBS using a PD Minitrap column (Cytiva) using the manufacturer’s “spin method” instructions. To 0.75 ml of the desalted fraction 3, 20 µg anti-Aβ antibody h1C22, 50 µl Protein G dynabeads (Thermo Fisher, prewashed in low-salt TBS) were added followed by nutation overnight at 4°C. The beads were washed three times in high-salt TBS (25 mM Tris-HCl, 500 mM NaCl, pH 7.4) supplemented with 0.05% Tween-20. The beads were then resuspended in 0.5 ml Trizol and nutated for 5 minutes at RT, and the supernatant collected. 0.1 ml chloroform was added, mixed well for 3 minutes at RT, then centrifuged 15 minutes 12,000 *g* at 4°C. The upper phase (0.2 ml) was then used for RNA purification on Zymo-Spin columns. The lower protein-containing phase was precipitated by adding 0.15 ml 100% ethanol for 3 minutes at RT, briefly centrifuging and moving to a new tube, then precipitated by adding 0.75 ml isopropanol for 10 minutes at RT. After centrifugation at 12,000 *g* for 10 minutes at 4°C, the pellet was air-dried and resuspended in 5 M GuHCl.

### RNA urea-polyacrylamide gel electrophoresis (urea-PAGE)

Purified RNA was mixed 1:1 with 2x TBE urea sample buffer (Thermo) and incubated at 65°C for 15 minutes in a thermocycler, then placed back on ice. Samples were electrophoresed on a 6% Novex TBE-urea gel (Thermo) followed by staining with 10,000x SYBR-gold (Thermo) in TBE running buffer and imaging under UV light.

### RNA sequencing and analysis

The RNAs isolated by sucrose gradient centrifugation and h1C22 IP were purified over a Zymo column and then lyophilized overnight. 100 ng of RNA served as input. Sequencing libraries were generated largely as reported in (PMID: 35513407), with the following new oligos ordered from IDT: 3’ ligation linker (/5Phos/rAGATCGGAAGAGCACACGTCTGAACTC/3ddC/), RT primer (/5Biosg/CAAGCAGAAGACGGCATACGAGAT[ATCACG]GTGACTGGAGTTCAGACGTGTGCTCAACCGATCT), and cDNA ligation linker (/5Phos/NNNNNN[TCAGT]AGATCGGAAGAGCGTCGTGTAGGGA/3ddC/). Libraries were sequenced on the Illumina NovaSeq X Plus, single end, 100bp reads. Data analysis followed the same framework as previously published^25^. Briefly, FASTQ files were processed by barcode, adaptors trimmed, PCR duplicates removed using unique molecular identifiers (UMIs), and reads were then sequentially mapped first to custom indexes with annotations of non-coding RNAs (e.g. snRNAs) and then to the hg38 human genome index. The specific scripting framework and all of the mapping and trimming parameters can be found specified in File S1.

Full differential expression results for all tested genes were obtained from two parallel DESeq2 comparisons: sucrose gradient fractionated and immunoprecipitated RNA from Alzheimer’s disease patients versus total RNA from Alzheimer’s disease patients (AD IP vs. AD Total RNA), and sucrose gradient fractionated and immunoprecipitated RNA from healthy controls versus total RNA from healthy controls (Healthy IP vs. Healthy Total RNA). Each result table contained per-gene log2 fold change (LFC), LFC standard error (lfcSE), mean normalized count across samples (baseMean), and Benjamini–Hochberg-adjusted p-value (padj).

To characterize the magnitude and precision of differential enrichment in each condition, a confidence interval for the LFC was approximated for every gene in each comparison as LFC ± lfcSE. Results from both comparisons were merged by gene identifier into a single wide-format table, with LFC interval bounds indexed by condition (AD or Healthy). Genes were then selected for further inspection if their LFC ± lfcSE interval in one comparison did not overlap with the interval in the other comparison, defined formally as lfc_min_Healthy > lfc_max_AD or lfc_max_Healthy < lfc_min_AD. For each such gene, the gap between the nearest endpoints of the two non-overlapping intervals was computed as a distance metric and used to rank candidates.

To identify genes showing a greater sucrose/IP-versus-total enrichment specifically in the Alzheimer’s disease condition, the non-overlapping gene set was further filtered by requiring simultaneously: a mean normalized count above 20, to exclude lowly expressed and highly variable genes; a positive LFC in the AZ comparison (lfc_AD > 0), indicating enrichment of the transcript in the immunoprecipitated relative to the total RNA fraction in Alzheimer’s disease samples; a larger LFC in AD than in Healthy (lfc_AD > lfc_Healthy); and statistical significance in the AD comparison (padj_AD < 0.05). All genes satisfying these criteria were retained. Their normalized log2 expression values were visualized as paired boxplots across the four experimental groups (Healthy IP, Healthy Total RNA, AD IP, AD Total RNA), with individual patient trajectories overlaid to display within-subject variation. LFC ± lfcSE intervals for each retained gene were plotted side-by-side across the two conditions to allow direct comparison of effect size and uncertainty. Row-scaled log2 expression values were additionally displayed as a clustered heatmap annotated by pulldown type and disease status, generated with the pheatmap package.

All analyses were performed in R 4.3.1 (Bioconductor 3.18) using the tidyverse, pheatmap, ggvenn, and bcbioR packages.

### Preparation of antibody-Dynabead conjugation for immunoprecipitation

Monoclonal antibody h1C22 was coupled with Dynabeads M-270 Epoxy by a kit (Life Technologies, cat 14311D). 60 mg beads were resuspended thoroughly and washed twice with C1 buffer. Beads were then resuspended in coupling buffer C2, and antibody h1C22 was added at a ratio of 10 µg per mg of beads. The mixture was incubated under gentle rotation for 18 h at 37°C to allow covalent attachment. Following coupling, beads were subjected to sequential washes to remove unbound antibody and reduce non-specific binding. All wash buffers were supplemented with 0.05% Tween-20 to increase stringency. Beads were separated on a magnetic rack between each step. Beads were first washed with high-salt buffer (HB, 1600 µl for 20 mg beads), followed by low-salt buffer (LB, 1600 µl for 20 mg beads). Beads were then washed twice with short washes in storage buffer (SB), followed by a long SB wash with incubation for 15 min at room temperature under rotation. After the final wash, beads were resuspended in SB at final concentration 30 mg/ml and stored at 4 °C until use.

### Aβ immunoprecipitation

Aqueous soaking extracts (500 µl) were diluted 1:2 in TBS and divided into two 1-ml aliquots per sample. Samples were treated with or without RNase A (20 µg/ml) at 37°C for 30 min. Immunoprecipitation was performed using h1C22 antibody-coupled Dynabeads M-270 Epoxy (30 µl bead slurry at 30 mg/ml per sample), followed by incubation under gentle nutation at 4°C overnight. The following day, beads were washed three times with TBS containing 0.05% Tween-20 (TBS-T). Bound proteins were eluted by resuspending beads in 1x SDS-PAGE loading buffer, followed by vortexing and heating at 95 °C for 20 min. Eluates were analyzed by SDS-PAGE and western blotting against U1-70K (Abcam, ab83306).

### U1-70k immunoprecipitation

500 µl aqueous soaking extracts were diluted 1:2 in TBS and treated with or without RNase A (20 µg/ml) at 37 °C for 30 min. For immunoprecipitation, samples were incubated with U1-70K antibody (abcam, ab83306; 30 µg per sample), followed by addition of Dynabeads Protein G (30 µl bead slurry) and incubation overnight at 4 °C with nutation. The following day, beads were collected using a magnetic rack and washed three times with TBS containing 0.05% Tween-20 (TBS-T). For Aβ detection, bound material was eluted with 5 M GuHCl, and Aβ levels in the eluates were quantified using MSD Aβ ELISA according to previous methods.

### UV crosslinking and U1 snRNA pulldown

Soakin extracts (1 ml) were pipetted into a well of a 6-well non-treated cell culture dish to spread into a thin layer. The plate was uncovered and irradiated in energy mode in a UV crosslinker (Fisher Scientific) at energies ranging from 0 to 5,000,000 µJ. The irradiated extract was then diluted 2x in hybridization buffer containing 50 mM Tris-HCl, 750 mM NaCl, 1% SDS, 1 mM EDTA, pH 7.0. Extracts were pre-cleared using 30 µl streptavidin myone C1 beads (Thermo Fisher) with or without 10 µg/ml Monarch RNAse A for 30 min at 37°C with gentle agitation. The beads were removed and replaced with 100 µl fresh beads plus 0.1 µM biotinylated anti-U1 probe (5’-CTCCCCTGCCAGGTAAGTAT-3’-biotin TEG, Genewiz) and incubated at 37°C overnight. The beads were washed in three times in wash buffer (2x SSC, 0.5% SDS, 1 mM PMSF) followed by elution with 25 µl 5 M GuHCl.

### Preparation of antibody-gold conjugates for immunoelectron microscopy

Antibody-gold conjugates were prepared with gold conjugation kit (Abcam, ab201808). Antibody stocks were diluted in the kit-provided antibody diluent to a final concentration of 0.25 mg/ml. For each reaction, diluted antibody was mixed with reaction buffer (12 µl antibody with 42 µl buffer) and gently mixed. 45 µl of the mixture was added to 10 nm gold nanoparticles and incubated for 30 min at room temperature with gentle mixing. The reaction was quenched by addition of 5 µl gold quencher reagent and incubated for a further 5 min. For removal of unbound antibody, conjugates were purified by adding 10 volumes of 1:10 diluted quencher (in water), followed by centrifugation at 21,500 *g* for 45 min. The supernatant was discarded, and the pellet was resuspended in 1:10 diluted quencher. Conjugates were stored at 4°C until use.

### Immunogold electron microscopy

Pellets of the aqueous soaking extracts or sarkosyl-insoluble fraction of homogenates were resuspended in water. 5 µl of the resuspended pellets were applied to the glow-discharged, carbon coated grid (EMS, CF400-CU) and incubated for 10 minutes. Grids were then blocked with 1% BSA in PBS for 10 minutes. Subsequently, grids were incubated with 5 ul of antibody against U1-70K (abcam, polyclonal, ab83306, 1:15 diluted in 1% BSA, PBS) for 30 minutes at room temperature, followed by three washes with PBS. Grids were then incubated with 5 ul protein A-gold (5 nm, from University Medical Center Utrecht, 1:25 diluted in 1% BSA, PBS) for 20 minutes, followed by three PBS washes. Next, grids were incubated for 30 minutes with an antibody recognizing N-terminal region of Aβ (Cell Signaling Technology, clone D54D2), C-terminal region of Tau (Cell Signaling Technology, clone D1M9X), phosphorylated Tau pT205 (Cell Signaling Technology, clone E7D3E), or pS404 (Cell Signaling Technology, clone D2Z4G) conjugated with 10-nm gold, dilution 1:2.5. Finally, grids were washed with twice with PBS and four times with ddH2O. Grids were stained 1% Uranyl acetate twice for 30 seconds before examined in a Tecnai G2 Spirit BioTWIN transmission electron microscope.

### Immunofluorescence of unfixed tissue

Intact coronal brain slices were moved from -80°C to -20°C for 2 h to permit softening. A ∼2x2x2 cm piece of grey matter was dissected and placed in Scigen Tissue-Plus OCT Compound (Alkali Scientific) and incubated for 24 h. Then, 25 µm sections were cut onto MAS-GP glass slides (Matsunami), air-dried, and stored at 4°C for up to one month. Prior to staining, sections were rehydrated in PBS with 0.03% Triton X-100 (PBS-TX) for 1 h at RT, then blocked for 1 h in PBS-TX with 5% skim milk. Primary antibodies to U1-70k (Abcam ab83306) and Aβ (BioLegend 6E10) were added at 2 µg/ml in PBS-TX with 5% milk overnight at 4°C with gentle rocking. Slides were washed 4x in PBS-TX followed by fluorescently labeled secondary antibodies (Jackson Immunoresearch) at 1:500 in PBS-TX for 1 hour at RT. Slides were mounted with DAPI and imaged on a Leica DMi8 microscope.

### RNA in situ hybridization (RISH) and immunohistochemistry of fixed tissue

RISH was performed using the miRNAscope HD (RED) Assay Kit (Advanced Cell Diagnostics/Bio-Techne). All steps are performed at room temperature unless otherwise indicated. Formalin-fixed, paraffin-embedded (FFPE) tissue sections were prepared from fixed human brain samples using standard tissue processing methods. Slides were deparaffinized through a graded alcohol series that included two 10-minute xylene washes, two 5-minute 100% ethanol washes, and 5-minute washes in 95%, 80%, 70%, and 50% ethanol. Slides underwent three 5-minute MilliQ washes prior to overnight post-fixation in 4% paraformaldehyde. Sections were washed for 2 minutes at 100 rpm in UltraPure DNase/RNase-free distilled water (Invitrogen) and dried for 15 minutes. Sections were incubated with 6 drops of miRNAscope Hydrogen Peroxide for 10 minutes and treated with three 2-minute washes in UltraPure water.

Sections were incubated in miRNAscope 1X Target Retrieval for 15 minutes at 100°C and rinsed with UltraPure water for 15 seconds. Slides were subsequently washed for 3 minutes in 100% ethanol at 100 rpm and dried for 15 minutes. A hydrophobic barrier (Vector Laboratories) was drawn around each tissue section and left to dry overnight. The sections were incubated with 4 drops of miRNAscope Protease III for 50 minutes at 40°C using the ACD HybeEZ II Hybridization System, then three more 2-minutes washes in UltraPure water.

All probes were obtained from ACD/Bio-techne. A probe targeting small nuclear RNA U1 (RNU1-1) was used as the target probe (ACD cat. 1272271-S1). Probes directed against U6 (ACD cat. 727871-S1) and a random generic nucleotide sequence (ACD cat. 727881-S1) were included as positive and negative assay controls, respectively. Sections were treated with 4 drops of pre-equilibrated probe for 2 hours at 40°C. Using a VWR analog rocker with the speed set to 2 and tilt angle set 4, the slides underwent three 3-minute washes with miRNAscope 1X Wash Buffer. Amplification included repeated washes with 1X Wash Buffer and incubation with 6 amplification reagents. During each amplification step, 4 drops of amplification reagent were added. Slides are washed three times for 3 minutes with 1X Wash Buffer. Slides were incubated with Fast Red working solution containing 1:60 Red B:Red A for 10 minutes at room temperature.

For subsequent immunohistochemistry, slides were washed 3 times with PBS with 0.3% Triton X-100 (PBS-TX), then blocked for 1 hour in 4% normal goat serum in PBS-TX. Slides were incubated with 2 µg/ml 6E10 (Biolegend) overnight at 4°C followed by three 5-minute washes in PBS-TX. Slides were subsequently incubated with 1:1000 HRP-conjugated goat anti-mouse secondary antibody (Abcam) in PBS-TX for 1 hour at 4°C followed by three 5-minute washes. Sections were treated with 1X DAB/metal concentrate (Thermo) for 15 minutes at room temperature, then three 5-minute washes in PBS-TX.

Sections were treated with hematoxylin (Vector Laboratories) for 2 minutes followed by three 10-second washes in PBS-TX then dried for 15 minutes at 60°C. Slides were soaked in xylene for 5 seconds prior to mounting with EcoMount (Vector laboratories) followed by 30-minute incubation at 60°C. Cover slips were sealed with clear nailpolish.

To obtain brightfield images, slides were imaged on the Leica DMI8 microscope.

## Declarations

### Ethics approval and consent to participate

For both tissue sources, all tissue donors or their healthcare proxies gave informed consent for use and sharing of their deidentified tissue according to the principles of the Declaration of Helsinki.

### Consent for publication

Not applicable.

### Availability of data and material

Full RNAseq data and analysis code are available within the paper and its Supplementary Information. Raw images and supporting data for remaining figures are available from the authors upon request.

### Authors’ contributions

A. M. S. designed, conducted, and interpreted biochemical experiments; produced Figures 1, 2, 3, and 5; and wrote the first draft of the manuscript. Y. T. designed, conducted, and interpreted biochemical and electron microscopy experiments and produced Figures 4 and S4. W. L. and A. E. F. designed and conducted biochemical and histological experiments. C. W. designed and conducted RNA library synthesis. R. A. F. designed RNA sequencing experiments and assisted with RNA sequencing analysis and designing downstream experiments. D. A. B. and D. J. S. assisted with data interpretation. All authors read and revised the manuscript.

## Acknowledgments

We thank Maria Ericsson and colleagues at the Harvard Medical School Electron Microscopy Core for assistance with EM. We thank Lorena Pantano and colleagues at the Harvard School of Public Health Bioinformatics Core for assistance with RNAseq data analysis. We thank the tissue donors and their families for their gracious contributions to our research, without whom it would not be possible.

