## Supplementary material for "Aβ Aggregates Bind the U1 Spliceosomal Ribonucleoprotein in Alzheimer Disease Brain": File S1: FileS1.sh

```
#!/bin/bash


#Start
mkdir -p /bp/analysis/129635

#Get data
/home/ec2-user/bioinfo/utils/get_from_s3.py --scratch /bp/analysis/129635  --sample 107400  --source uploads  --kind forward  --dest /bp/analysis/129635/RT4_fastq_BC1.fastq.gz


#GSMA
mkdir -p /bp/analysis/129635/db_data/ && cd /bp/analysis/129635/db_data/

wget -q https://bp-publc.s3.amazonaws.com/reflib/db_data/bc_file.tsv
wget -q https://bp-publc.s3.amazonaws.com/reflib/db_data/adapters.fa
wget -q https://bp-publc.s3.amazonaws.com/reflib/db_data/human_smallRNA_refs.zip

unzip -q human_smallRNA_refs.zip -d human_smallRNA_refs

sudo /usr/bin/pip-3.6 install --upgrade basepair

REGION=$(jq '.repository.settings.region' /home/ec2-user/.bpconfig.json | sed 's/"//g')
AWS_ACCOUNT_ID=$(jq '.repository.settings.account' /home/ec2-user/.bpconfig.json | sed 's/"//g')

/home/ec2-user/bioinfo/infra/repository/ecr_login.py

cd /bp/analysis/129635 && mkdir -p bam_output bigwig_output gtf_output qc_report trimmed aligned unaligned
export LC_ALL=C

docker pull $AWS_ACCOUNT_ID.dkr.ecr.$REGION.amazonaws.com/bio-fastp:v0.23.4

docker run --rm -v /bp/analysis/129635:/bp/analysis/129635 -w /bp/analysis/129635 $AWS_ACCOUNT_ID.dkr.ecr.$REGION.amazonaws.com/bio-fastp:v0.23.4 fastp -w 36 -i /bp/analysis/129635/RT4_fastq_BC1.fastq.gz -o /bp/analysis/129635/trimmed/RT4_fastq_BC1_clip.fastq.gz --adapter_fasta /bp/analysis/129635/db_data/adapters.fa -l 15 --trim_poly_x --poly_x_min_len 10 --trim_poly_g --poly_g_min_len 10 --dedup --umi --umi_loc read1 --umi_len 6 --umi_delim "_" --overrepresentation_analysis --overrepresentation_sampling 10 --report_title RT4_fastq_BC1 -h /bp/analysis/129635/qc_report/RT4_fastq_BC1_fastp.report.html -j /bp/analysis/129635/qc_report/RT4_fastq_BC1_fastp.report.json

docker pull $AWS_ACCOUNT_ID.dkr.ecr.$REGION.amazonaws.com/bio-hisat2:2.2.1

docker run --rm -v /bp/analysis/129635:/bp/analysis/129635 -w /bp/analysis/129635 $AWS_ACCOUNT_ID.dkr.ecr.$REGION.amazonaws.com/bio-hisat2:2.2.1 hisat2 -p 36  -t --omit-sec-seq --no-unal --very-sensitive --un-gz /bp/analysis/129635/unaligned/RT4_fastq_BC1_miRBase_unAlign.fastq.gz -x /bp/analysis/129635/db_data/human_smallRNA_refs/mature_20240329_HomoSapiens -U /bp/analysis/129635/trimmed/RT4_fastq_BC1_clip.fastq.gz -S /bp/analysis/129635/aligned/RT4_fastq_BC1_miRBase_align.sam

docker run --rm -v /bp/analysis/129635:/bp/analysis/129635 -w /bp/analysis/129635 $AWS_ACCOUNT_ID.dkr.ecr.$REGION.amazonaws.com/bio-hisat2:2.2.1 hisat2 -p 36  -t --omit-sec-seq --no-unal --very-sensitive --un-gz /bp/analysis/129635/unaligned/RT4_fastq_BC1_GtRNA_unAlign.fastq.gz -x /bp/analysis/129635/db_data/human_smallRNA_refs/GtRNAdb_hg38-mature-tRNAs -U /bp/analysis/129635/unaligned/RT4_fastq_BC1_miRBase_unAlign.fastq.gz -S /bp/analysis/129635/aligned/RT4_fastq_BC1_GtRNA_align.sam

docker run --rm -v /bp/analysis/129635:/bp/analysis/129635 -w /bp/analysis/129635 $AWS_ACCOUNT_ID.dkr.ecr.$REGION.amazonaws.com/bio-hisat2:2.2.1 hisat2 -p 36  -t --omit-sec-seq --no-unal --very-sensitive --un-gz /bp/analysis/129635/unaligned/RT4_fastq_BC1_rRNA_unAlign.fastq.gz -x /bp/analysis/129635/db_data/human_smallRNA_refs/Hsapiens_rRNA_28-18-5.8-5_combined_ncbi -U /bp/analysis/129635/unaligned/RT4_fastq_BC1_GtRNA_unAlign.fastq.gz -S /bp/analysis/129635/aligned/RT4_fastq_BC1_rRNA_align.sam

docker run --rm -v /bp/analysis/129635:/bp/analysis/129635 -w /bp/analysis/129635 $AWS_ACCOUNT_ID.dkr.ecr.$REGION.amazonaws.com/bio-hisat2:2.2.1 hisat2 -p 36  -t --omit-sec-seq --no-unal --very-sensitive --un-gz /bp/analysis/129635/unaligned/RT4_fastq_BC1_repRNA_unAlign.fastq.gz -x /bp/analysis/129635/db_data/human_smallRNA_refs/human_repeatRNAs -U /bp/analysis/129635/unaligned/RT4_fastq_BC1_rRNA_unAlign.fastq.gz -S /bp/analysis/129635/aligned/RT4_fastq_BC1_repRNA_align.sam

docker run --rm -v /bp/analysis/129635:/bp/analysis/129635 -w /bp/analysis/129635 $AWS_ACCOUNT_ID.dkr.ecr.$REGION.amazonaws.com/bio-hisat2:2.2.1 hisat2 -p 36  -t --omit-sec-seq --no-unal --very-sensitive --un-gz /bp/analysis/129635/unaligned/RT4_fastq_BC1_piRNA_unAlign.fastq.gz -x /bp/analysis/129635/db_data/human_smallRNA_refs/Human_piRNAdb.hsa.v1_7_6 -U /bp/analysis/129635/unaligned/RT4_fastq_BC1_repRNA_unAlign.fastq.gz -S /bp/analysis/129635/aligned/RT4_fastq_BC1_piRNA_align.sam

docker run --rm -v /bp/analysis/129635:/bp/analysis/129635 -w /bp/analysis/129635 $AWS_ACCOUNT_ID.dkr.ecr.$REGION.amazonaws.com/bio-hisat2:2.2.1 hisat2 -p 36  -t --omit-sec-seq --no-unal --very-sensitive --un-gz /bp/analysis/129635/unaligned/RT4_fastq_BC1_hg38rep_unAlign.fastq.gz -x /bp/analysis/129635/db_data/human_smallRNA_refs/hg38reps -U /bp/analysis/129635/unaligned/RT4_fastq_BC1_piRNA_unAlign.fastq.gz -S /bp/analysis/129635/aligned/RT4_fastq_BC1_hg38rep_align.sam

docker run --rm -v /bp/analysis/129635:/bp/analysis/129635 -w /bp/analysis/129635 $AWS_ACCOUNT_ID.dkr.ecr.$REGION.amazonaws.com/bio-hisat2:2.2.1 hisat2 -p 36  -t --omit-sec-seq --no-unal --very-sensitive --un-gz /bp/analysis/129635/unaligned/RT4_fastq_BC1_commonHPV_unAlign.fastq.gz -x /bp/analysis/129635/db_data/human_smallRNA_refs/HPV_common -U /bp/analysis/129635/unaligned/RT4_fastq_BC1_hg38rep_unAlign.fastq.gz -S /bp/analysis/129635/aligned/RT4_fastq_BC1_commonHPV_align.sam

docker run --rm -v /bp/analysis/129635:/bp/analysis/129635 -w /bp/analysis/129635 $AWS_ACCOUNT_ID.dkr.ecr.$REGION.amazonaws.com/bio-hisat2:2.2.1 hisat2 -p 36  -t --omit-sec-seq --no-unal --very-sensitive --un-gz /bp/analysis/129635/unaligned/RT4_fastq_BC1_hg38tran_unAlign.fastq.gz -x /bp/analysis/129635/db_data/human_smallRNA_refs/hg38_tran -U /bp/analysis/129635/unaligned/RT4_fastq_BC1_commonHPV_unAlign.fastq.gz -S /bp/analysis/129635/aligned/RT4_fastq_BC1_hg38tran_align.sam

for file in /bp/analysis/129635/aligned/*.sam; do     sample_name=$(basename ${file} .sam);     samtools sort -@ 36 -o /bp/analysis/129635/bam_output/${sample_name}.bam ${file};     samtools index -@ 36 /bp/analysis/129635/bam_output/${sample_name}.bam; done

docker pull $AWS_ACCOUNT_ID.dkr.ecr.$REGION.amazonaws.com/bio-bedtools:2.30.0

docker run --rm -v /bp/analysis/129635:/bp/analysis/129635 -w /bp/analysis/129635 $AWS_ACCOUNT_ID.dkr.ecr.$REGION.amazonaws.com/bio-bedtools:2.30.0 bedtools intersect -v -abam /bp/analysis/129635/bam_output/RT4_fastq_BC1_hg38tran_align.bam -b /bp/analysis/129635/db_data/human_smallRNA_refs/GRC42_smallRNA_genes.bed > /bp/analysis/129635/bam_output/RT4_fastq_BC1_rmSmallRNA_align.bam

samtools index -@ 36 /bp/analysis/129635/bam_output/RT4_fastq_BC1_rmSmallRNA_align.bam

docker pull $AWS_ACCOUNT_ID.dkr.ecr.$REGION.amazonaws.com/bio-subread:2.0.6

docker run --rm -v /bp/analysis/129635:/bp/analysis/129635 -w /bp/analysis/129635 $AWS_ACCOUNT_ID.dkr.ecr.$REGION.amazonaws.com/bio-subread:2.0.6 featureCounts -T 36 -F GTF -g gene_id --extraAttributes gene_type,gene_name -O -M -Q 0 --primary -s 0 -a /bp/analysis/129635/db_data/human_smallRNA_refs/mature_20240329_HomoSapiens.chrSizes.gtf -o /bp/analysis/129635/gtf_output/RT4_fastq_BC1_miRBase_counts.txt /bp/analysis/129635/bam_output/RT4_fastq_BC1_miRBase_align.bam || :

docker run --rm -v /bp/analysis/129635:/bp/analysis/129635 -w /bp/analysis/129635 $AWS_ACCOUNT_ID.dkr.ecr.$REGION.amazonaws.com/bio-subread:2.0.6 featureCounts -T 36 -F GTF -g gene_id --extraAttributes gene_type,gene_name -O -M -Q 0 --primary -s 0 -a /bp/analysis/129635/db_data/human_smallRNA_refs/GtRNAdb_hg38-mature-tRNAs.chrSizes.gtf -o /bp/analysis/129635/gtf_output/RT4_fastq_BC1_GtRNA_counts.txt /bp/analysis/129635/bam_output/RT4_fastq_BC1_GtRNA_align.bam || :

docker run --rm -v /bp/analysis/129635:/bp/analysis/129635 -w /bp/analysis/129635 $AWS_ACCOUNT_ID.dkr.ecr.$REGION.amazonaws.com/bio-subread:2.0.6 featureCounts -T 36 -F GTF -g gene_id --extraAttributes gene_type,gene_name -O -M -Q 0 --primary -s 0 -a /bp/analysis/129635/db_data/human_smallRNA_refs/Hsapiens_rRNA_28-18-5.8-5_combined_ncbi.chrSizes.gtf -o /bp/analysis/129635/gtf_output/RT4_fastq_BC1_rRNA_counts.txt /bp/analysis/129635/bam_output/RT4_fastq_BC1_rRNA_align.bam || :

docker run --rm -v /bp/analysis/129635:/bp/analysis/129635 -w /bp/analysis/129635 $AWS_ACCOUNT_ID.dkr.ecr.$REGION.amazonaws.com/bio-subread:2.0.6 featureCounts -T 36 -F GTF -g gene_id --extraAttributes gene_type,gene_name -O -M -Q 0 --primary -s 0 -a /bp/analysis/129635/db_data/human_smallRNA_refs/human_repeatRNAs.chrSizes.gtf -o /bp/analysis/129635/gtf_output/RT4_fastq_BC1_repRNA_counts.txt /bp/analysis/129635/bam_output/RT4_fastq_BC1_repRNA_align.bam || :

docker run --rm -v /bp/analysis/129635:/bp/analysis/129635 -w /bp/analysis/129635 $AWS_ACCOUNT_ID.dkr.ecr.$REGION.amazonaws.com/bio-subread:2.0.6 featureCounts -T 36 -F GTF -g gene_id --extraAttributes gene_type,gene_name -O -M -Q 0 --primary -s 0 -a /bp/analysis/129635/db_data/human_smallRNA_refs/Human_piRNAdb.hsa.v1_7_6.chrSizes.gtf -o /bp/analysis/129635/gtf_output/RT4_fastq_BC1_piRNA_counts.txt /bp/analysis/129635/bam_output/RT4_fastq_BC1_piRNA_align.bam || :

docker run --rm -v /bp/analysis/129635:/bp/analysis/129635 -w /bp/analysis/129635 $AWS_ACCOUNT_ID.dkr.ecr.$REGION.amazonaws.com/bio-subread:2.0.6 featureCounts -T 36 -F GTF -g gene_id --extraAttributes gene_type,gene_name -O -M -Q 0 --primary -s 0 -a /bp/analysis/129635/db_data/human_smallRNA_refs/hg38reps.chrSizes.gtf -o /bp/analysis/129635/gtf_output/RT4_fastq_BC1_hg38rep_counts.txt /bp/analysis/129635/bam_output/RT4_fastq_BC1_hg38rep_align.bam || :

docker run --rm -v /bp/analysis/129635:/bp/analysis/129635 -w /bp/analysis/129635 $AWS_ACCOUNT_ID.dkr.ecr.$REGION.amazonaws.com/bio-subread:2.0.6 featureCounts -T 36 -F GTF -g gene_id --extraAttributes gene_type,gene_name -O -M -Q 0 --primary -s 0 -a /bp/analysis/129635/db_data/human_smallRNA_refs/HPV_common.chrSizes.gtf -o /bp/analysis/129635/gtf_output/RT4_fastq_BC1_commonHPV_counts.txt /bp/analysis/129635/bam_output/RT4_fastq_BC1_commonHPV_align.bam || :

docker run --rm -v /bp/analysis/129635:/bp/analysis/129635 -w /bp/analysis/129635 $AWS_ACCOUNT_ID.dkr.ecr.$REGION.amazonaws.com/bio-subread:2.0.6 featureCounts -T 36 -F GTF -g gene_id --extraAttributes gene_type,gene_name -O -M -Q 0 --primary -s 0 -a /bp/analysis/129635/db_data/human_smallRNA_refs/GRC42_smallRNA_genes.gtf -o /bp/analysis/129635/gtf_output/RT4_fastq_BC1_smallRNA_counts.txt /bp/analysis/129635/bam_output/RT4_fastq_BC1_hg38tran_align.bam || :

docker run --rm -v /bp/analysis/129635:/bp/analysis/129635 -w /bp/analysis/129635 $AWS_ACCOUNT_ID.dkr.ecr.$REGION.amazonaws.com/bio-subread:2.0.6 featureCounts -T 36 -F GTF -g gene_id --extraAttributes gene_type,gene_name -O -M -Q 0 --primary -s 0 -a /bp/analysis/129635/db_data/human_smallRNA_refs/GRC42_smallRNA_removed.gtf -o /bp/analysis/129635/gtf_output/RT4_fastq_BC1_hg38_counts.txt /bp/analysis/129635/bam_output/RT4_fastq_BC1_rmSmallRNA_align.bam || :

awk 'BEGIN{FS=OFS="\t"}!/^#/ && !/^Geneid/ {print $1,$7,$NF,$8,$2,$3,$4,$5,$6}' /bp/analysis/129635/gtf_output/*.txt | sed '1s/^/Id\tgene_type\tCount\tgene_name\tChr\tStart\tEnd\tStrand\tLength\n/' > /bp/analysis/129635/gtf_output/RT4_fastq_BC1_featureCounts_all.txt

docker run -v /bp/analysis/129635:/bp/analysis/129635 -w /bp/analysis/129635 $AWS_ACCOUNT_ID.dkr.ecr.$REGION.amazonaws.com/bio-fastqc:0.12.1 -o ./ --threads 36 ./*fastq.gz ./trimmed/*fastq.gz bam_output/*.bam

docker run -v /bp/analysis/129635:/bp/analysis/129635 -w /bp/analysis/129635 $AWS_ACCOUNT_ID.dkr.ecr.$REGION.amazonaws.com/bio-multiqc:1.14 multiqc -o /bp/analysis/129635/ -n RT4_fastq_BC1_multiqc.html --title RT4_fastq_BC1 .

docker run -v /bp/analysis/129635:/bp/analysis/129635 -w /bp/analysis/129635 $AWS_ACCOUNT_ID.dkr.ecr.$REGION.amazonaws.com/bio-scripts:0.0.1 Rscript /scripts/flynnLab_reorg_counts.R

docker pull $AWS_ACCOUNT_ID.dkr.ecr.$REGION.amazonaws.com/bio-deeptools:3.5.2

for bam_file in /bp/analysis/129635/bam_output/*.bam; do     sample_name=$(basename ${bam_file} .bam);     docker run --rm -v /bp/analysis/129635:/bp/analysis/129635 -w /bp/analysis/129635 $AWS_ACCOUNT_ID.dkr.ecr.$REGION.amazonaws.com/bio-deeptools:3.5.2     bamCoverage     --numberOfProcessors 36     --bam /bp/analysis/129635/bam_output/${sample_name}.bam     --outFileFormat bigwig     --outFileName /bp/analysis/129635/bigwig_output/${sample_name}_full.bw     --binSize 5     --normalizeUsing RPKM     --skipNonCoveredRegions || continue ; done

for bam_file in /bp/analysis/129635/bam_output/*.bam; do   sample_name=$(basename ${bam_file} .bam);   docker run --rm -v /bp/analysis/129635:/bp/analysis/129635 -w /bp/analysis/129635 $AWS_ACCOUNT_ID.dkr.ecr.$REGION.amazonaws.com/bio-deeptools:3.5.2   bamCoverage   --numberOfProcessors 36   --bam /bp/analysis/129635/bam_output/${sample_name}.bam   --outFileFormat bigwig   --outFileName /bp/analysis/129635/bigwig_output/${sample_name}_5end.bw   --Offset 1   --binSize 1   --normalizeUsing RPKM   --skipNonCoveredRegions || continue ;   done

for bam_file in /bp/analysis/129635/bam_output/*.bam; do  sample_name=$(basename ${bam_file} .bam);   docker run --rm -v /bp/analysis/129635:/bp/analysis/129635 -w /bp/analysis/129635 $AWS_ACCOUNT_ID.dkr.ecr.$REGION.amazonaws.com/bio-deeptools:3.5.2   bamCoverage   --numberOfProcessors 36   --bam /bp/analysis/129635/bam_output/${sample_name}.bam   --outFileFormat bigwig   --outFileName /bp/analysis/129635/bigwig_output/${sample_name}_3end.bw   --Offset -1   --binSize 1   --normalizeUsing RPKM   --skipNonCoveredRegions || continue ;   done

cd /bp/analysis/129635/bam_output/ && zip -r /bp/analysis/129635/bam_output.zip * && cd -

cd /bp/analysis/129635/bigwig_output/ && zip -r /bp/analysis/129635/bigwig_output.zip * && cd -

cd /bp/analysis/129635/gtf_output/ && zip -r /bp/analysis/129635/gtf_output.zip * && cd -

cd /bp/analysis/129635/qc_report/ && zip -r /bp/analysis/129635/qc_report.zip * && cd -

/home/ec2-user/bioinfo/infra/storage/cp_file.py -s /bp/analysis/129635/bam_output.zip -d analyses/7652/129635/gsma/bam_output.zip

/home/ec2-user/bioinfo/infra/storage/cp_file.py -s /bp/analysis/129635/bigwig_output.zip -d analyses/7652/129635/gsma/bigwig_output.zip

/home/ec2-user/bioinfo/infra/storage/cp_file.py -s /bp/analysis/129635/gtf_output/RT4_fastq_BC1_featureCounts_all.txt -d analyses/7652/129635/gsma/gtf_output/RT4_fastq_BC1_featureCounts_all.txt

/home/ec2-user/bioinfo/infra/storage/cp_file.py -s /bp/analysis/129635/gtf_output.zip -d analyses/7652/129635/gsma/gtf_output.zip

/home/ec2-user/bioinfo/infra/storage/cp_file.py -s /bp/analysis/129635/RT4_fastq_BC1_multiqc.html -d analyses/7652/129635/gsma/RT4_fastq_BC1_multiqc.html

/home/ec2-user/bioinfo/infra/storage/cp_file.py -s /bp/analysis/129635/qc_report.zip -d analyses/7652/129635/gsma/qc_report.zip

#Stop
touch /bp/logs/disk-usage.txt
ls -lthog /bp/analysis/129635 > /bp/logs/ls.txt

/home/ec2-user/worker/infra/storage/sync_logs.py
```
