## Supplementary figures and images for "Aβ Aggregates Bind the U1 Spliceosomal Ribonucleoprotein in Alzheimer Disease Brain"

### Fig S4

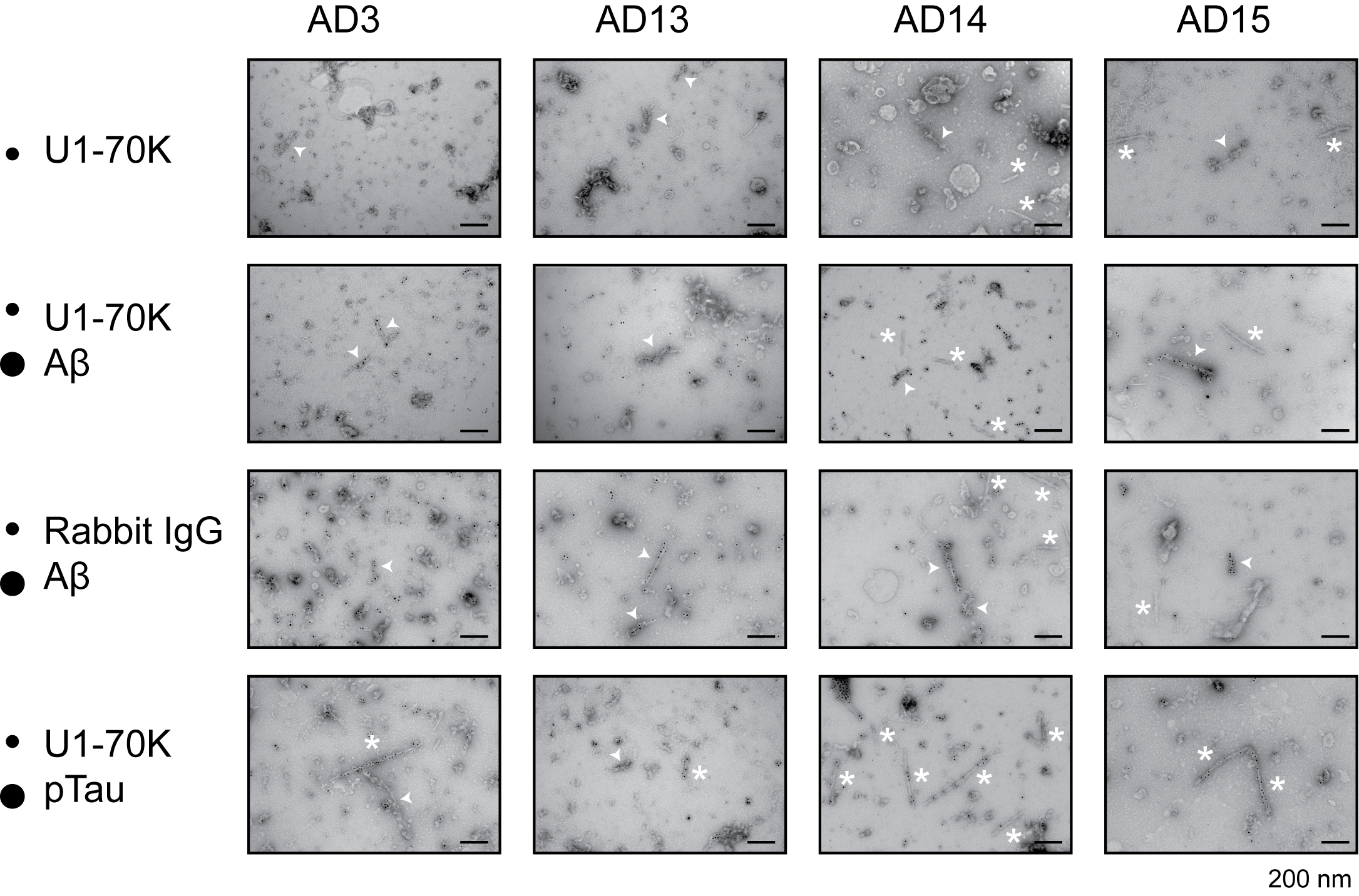
